# The position of single-base deletions in the VNTR sequence of the carboxyl ester lipase (*CEL*) gene determines pathogenicity

**DOI:** 10.1101/2020.12.23.424204

**Authors:** Anny Gravdal, Xunjun Xiao, Miriam Cnop, Khadija El Jellas, Pål R. Njølstad, Mark E. Lowe, Bente B. Johansson, Anders Molven, Karianne Fjeld

**Author notes:** To whom correspondence should be addressed: The Gade Laboratory for Pathology, Department of Clinical Medicine, University of Bergen, Jonas Lies vei 87, N-5021 Bergen, Norway.

## Abstract

Variable number of tandem repeat (VNTR) sequences present in the genome can have functional consequences that contribute to human disease. This is the case for the *CEL* gene, which encodes the digestive enzyme carboxyl ester lipase. *CEL* has a VNTR located in exon 11, and rare single-base deletions (DELs) within this region cause MODY8, an inherited disorder characterized by exocrine pancreatic dysfunction and diabetes. Here, we have studied how the position of single-base deletions within the *CEL* VNTR affects the protein’s pathogenic properties. We investigated four naturally occurring *CEL* variants with single-base deletions in different VNTR segments (DEL1, DEL4, DEL9, DEL13), of which only DEL1 and DEL4 have been observed in MODY8 patients. When expressed in a cellular model system, only DEL1 and DEL4 exhibited significantly reduced secretion and increased intracellular aggregation compared to normal CEL. We found that all DEL variants had slightly decreased enzymatic activity and that their level of O-glycosylation was affected. Moreover, only DEL1 and DEL4 significantly increased endoplasmic reticulum (ER) stress. In conclusion, *CEL* single-base deletion variants have the highest pathogenic potential when the mutational event has taken place in the proximal VNTR part, resulting in the longest aberrant protein tails. Thus, DEL1 and DEL4 are pathogenic *CEL* variants, whereas we consider DEL13 as benign and DEL9 as likely benign. These findings have implications for our understanding of how *CEL* mutations cause pancreatic disease through protein misfolding and proteotoxicity, leading to ER stress and activation of the unfolded protein response.

---

Variable number of tandem repeat (VNTR) is a term used to describe short DNA sequence motifs that are consecutively repeated several times in the genome (1). As these motifs are very polymorphic and inherited in a Mendelian pattern, they have had a tremendous impact on our ability to identify individuals based on biological samples (2). The VNTRs are most often located in non-coding regions of the genome, but some are found in promoter regions (3) or within coding DNA sequences (4). In such cases, a VNTR may influence the expression levels or functional properties of the corresponding protein. Moreover, as repeated DNA motifs are prone to undergo mutations by DNA slippage and unequal crossing-over, VNTRs affecting the protein product are intriguing candidates for human disease associations (1).

One such example is found in the *CEL* gene, encoding the digestive enzyme carboxyl ester lipase, which is mainly expressed in the acinar cells of the pancreas (5). *CEL* contains 11 exons, where the last exon has a VNTR that consists of nearly identical 33-base pair segments, each encoding 11 amino acids (5). The number of repeated segments within the VNTR may vary between three and 23. However, in all populations examined so far, the most common *CEL* allele contains 16 repeats, encoding a protein with a predicted molecular mass of 79 kDa (6–11) (**Fig. 1**). As the CEL protein is both N- and O-glycosylated, the observed mass is considerably larger (12,13). In particular, threonine residues within the VNTR-encoded tail region undergo O-glycosylation (13). This modification may serve to prevent rapid degradation of the CEL protein, but might also facilitate secretion or increase solubility (14). Intriguingly, the pattern of O-glycosylation in CEL reflects the ABO blood group of the individual (15).

**Figure 1.**
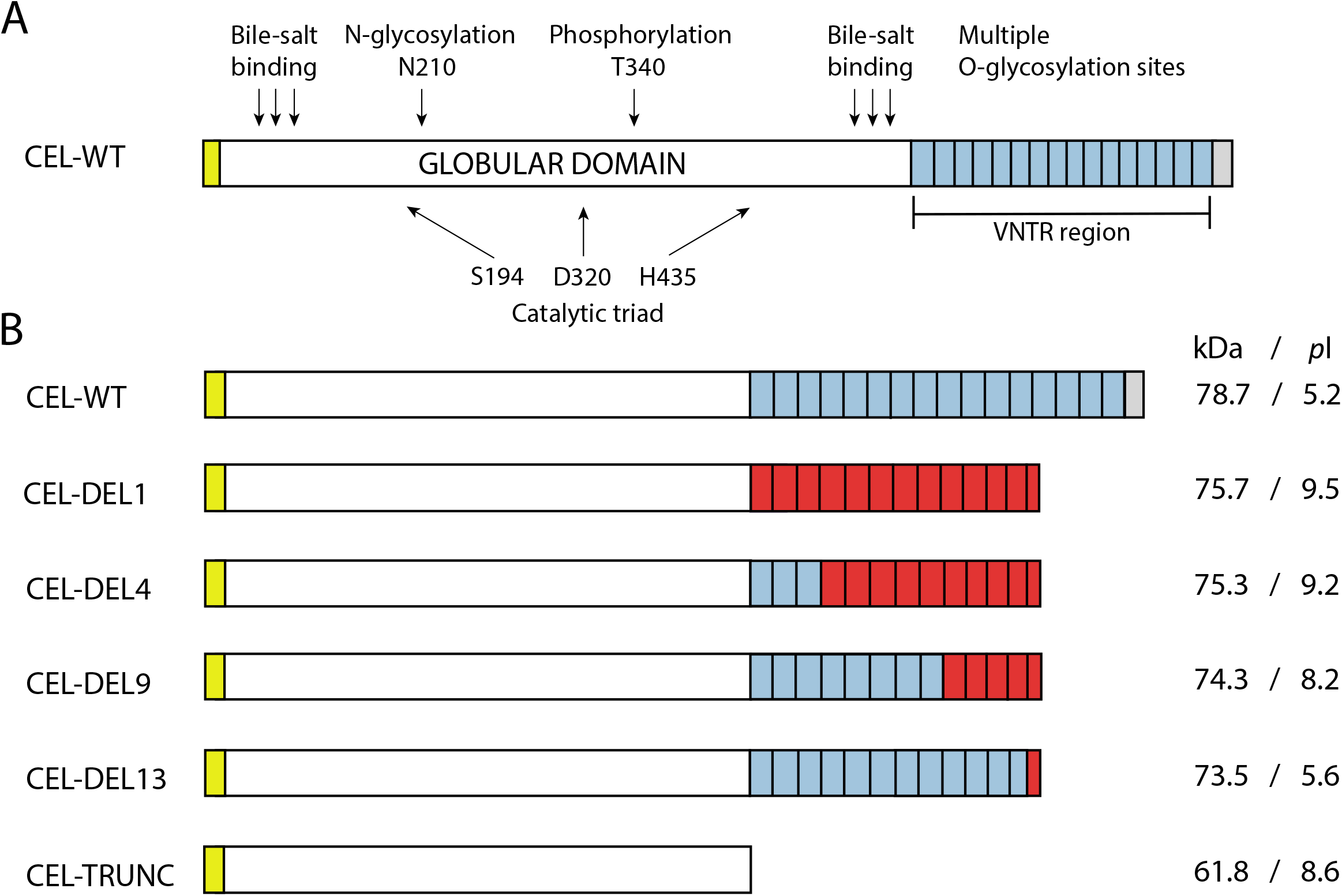
Schematic overview of the different CEL protein variants encoded by constructs used in the present study. *A*, the normal CEL protein (CEL-WT) with 16 repeats, each consisting of 11 amino acids and encoded by the VNTR. This is the most common CEL variant in the general population. The yellow box symbolizes the N-terminal signal peptide, whereas light blue boxes represent the normal repeats. The unique C-terminal sequence of CEL-WT is symbolized by a grey box. Also indicated are residues required for bile salt binding, N- and O-glycosylation, phosphorylation, and catalysis. *B*, overview of the different CEL protein variants investigated. Predicted molecular mass (kDa) and the isoelectric point (*pI*) of each variant are shown on the right side. All deletion constructs were based on a 16-repeat backbone. Light blue boxes represent normal repeat sequences, whereas red boxes indicate aberrant repeats resulting from the frameshifts introduced by singe-base pair deletions in the *CEL* VNTR. CEL-TRUNC is an artificial variant lacking the VNTR region and was included as control construct. The elements of A and B are not drawn to scale.

A single-base deletion in the first repeat of the *CEL* VNTR leads to MODY8, a dominantly inherited syndrome of exocrine pancreatic dysfunction and diabetes (16). The corresponding pathogenic protein has been referred to as CEL-MODY or CEL-MUT in previous publications (17,18) but will hereafter be denoted CEL-DEL1 (abbreviated: DEL1) to emphasize that the deletion is positioned in the first repeat. Another single-base deletion in the fourth repeat of the *CEL* VNTR was found in a family with a similar phenotype as the original MODY8 family (16), and this protein will be denoted DEL4. Due to frameshifts and premature stop codons, DEL1 and DEL4 are both truncated CEL variants with aberrant protein C-termini (Fig. 1). Very rare single-base deletions in the ninth and thirteenth repeat of the *CEL* VNTR (encoding protein variants DEL9 and DEL13, respectively; Fig. 1) have been observed in healthy controls (unpublished data). It is unknown whether these variants also associate with disease.

We have previously shown that DEL1 has a high propensity to form both intra- and extracellular aggregates (18). We have also found that after secretion, there is cellular reuptake followed by lysosomal degradation of the DEL1 protein, resulting in reduced viability of pancreatic cell lines (17,19). Furthermore, we have reported that DEL1 causes endoplasmic reticulum (ER) stress, induction of the unfolded protein response and subsequent apoptosis (20). Taken together, these results indicate that the CEL protein in MODY8 has acquired a negative gain-of-function effect, and that the disease follows the protein misfolding-dependent pathway of pancreatitis (21).

The aim of the present study was to examine the pathogenicity of single-base deletions naturally occurring in the *CEL* VNTR. We tested how the position of a deletion within this region affects the functional properties of the encoded CEL protein variant. We therefore compared the DEL1 and DEL4 variants causing MODY8 with two deletions that we have observed in the general population (DEL9, DEL13). Our data suggest a correlation between the position of a *CEL* VNTR single-base deletion and its functional impact on the protein, where increasingly distal mutations have less pathogenic potential.

## RESULTS

### Cellular properties of CEL proteins are influenced by the V5/His tag

In previous cellular studies of the pathogenic DEL1 variant, we have expressed the protein both with and without a V5/His epitope tag fused to the C-terminus (15–20). While epitope tags are very useful tools for protein purification and detection, the charged and hydrophilic residues of such tags may sometimes have a significant impact on the biochemical properties of the protein (22,23). Before initiating the functional study of our set of CEL deletion variants, we therefore investigated how the V5/His tag might influence the CEL protein.

HEK293 cells were transiently transfected with plasmids expressing DEL1 with and without tag, followed by either cellular fractionation and western blotting or immunocytochemistry. As controls, we included a construct encoding the normal CEL protein with 16 repeats (wildtype, WT) and a construct with a stop codon introduced at the beginning of VNTR segment 1, encoding a truncated CEL protein (TRUNC, Fig. 1). We found the WT and DEL1 proteins expressed without the V5/His tag to be less secreted than the tagged variants (Supporting Fig. 1*A*). In addition, the untagged variants showed slightly stronger bands in the insoluble pellet fraction, indicating that they are more prone to aggregate. Consistent with these findings, confocal imaging demonstrated a stronger intracellular signal for the untagged WT and DEL1 proteins than for their tagged counterparts (Supporting Fig. 1*B*). The TRUNC protein did not display visible differences with or without the tag in our experiments.

Based on these results, we concluded that the V5/His tag may increase the solubility of CEL protein variants, possibly via interactions involving the VNTR-encoded region. In previous studies with tagged variants (17–19), we may therefore have underestimated the detrimental cellular effects of DEL1. Consequently, we decided to use only untagged protein variants in the present study.

### VNTR length and composition affects CEL protein secretion and intracellular distribution

We have previously shown that the disease-causing DEL1 variant exhibits intracellular retention and reduced secretion (20). To test how the position of a single-base deletion within the VNTR may affect secretion and intracellular distribution of the protein, we compared four different CEL variants harboring single-base deletions in the first, fourth, ninth and thirteenth repeat, respectively (DEL1, DEL4, DEL9, DEL13; Fig. 1). The WT protein and the TRUNC variant were included as references. HEK293 cells were transiently transfected with the six constructs, and the medium, soluble lysate and detergent-insoluble pellet were analyzed by western blotting.

When compared with CEL-WT, secretion and intracellular distribution varied substantially between the variants (**Fig. 2**). We found significantly reduced secretion for DEL1 (*p* < 0.0001), DEL4 (*p* = 0.0045) and TRUNC (*p* < 0.001). In contrast, variants DEL9 and DEL13 displayed a secretion level similar to that of the WT protein. In the pellet fraction, we observed stronger signals for DEL1 (*p* = 0.017), DEL4 (*p* = 0.045) and TRUNC (*p* = 0.039). DEL13 behaved in the same way as the WT protein. The variant DEL9 displayed an apparently stronger band intensity than CEL-WT; however, the difference was not statistically significant. Moreover, significantly lower levels of all variants (DEL1: *p* = 0.002, DEL4: *p* = 0.004, DEL9: *p* = 0.007, DEL 13: *p* = 0.049) were detected in the soluble lysate fraction (Fig. 2).

**Figure 2.**
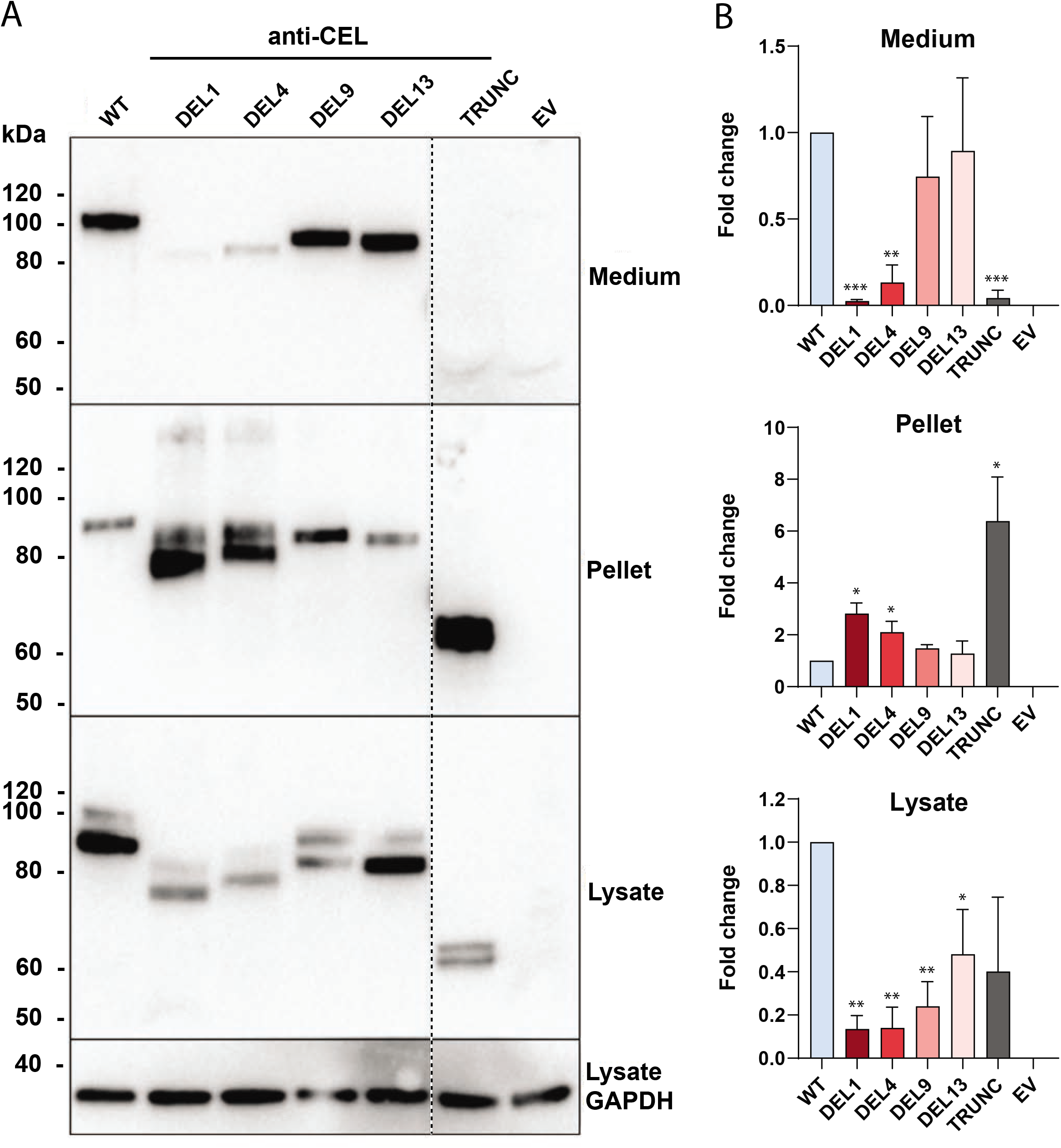
Secretion and cellular distribution of different CEL protein variants. *A*, HEK293 cells were transiently transfected with plasmids encoding CEL-WT, the four deletion variants (DEL1, DEL4, DEL9, DEL13) or CEL TRUNC, and subjected to cellular fractionation and western blotting. Cells transfected with empty vector (EV) were used as negative control, whereas GAPDH expression was monitored for loading control. Secretion was assessed as CEL levels detected in the conditioned medium. Intracellular distribution was evaluated by analyzing the soluble (lysate) and insoluble (pellet) fractions after lysis of the cells. The stippled line indicates an unrelated lane removed from the images. One representative image of three independent experiments is shown. *B*, quantification of band intensity of three individual western blot images, adjusted to the GAPDH levels and normalized to the CEL-WT lane. Error bars are standard deviation (SD). Statistically significant differences compared with CEL-WT are indicated as follows: *, p < 0.05; **, p < 0.01; ***, p < 0.001.

### The extent of O-glycosylation varies between CEL deletion variants

Interestingly, the four deletion variants migrated with clearly different band sizes in the SDS-PAGE gels despite having very similar predicted molecular masses (compare Fig. 1 and Fig. 2). This suggested that the variants differ at the level of post-translational modification. Since CEL is heavily O-glycosylated (13,15,24), a likely explanation for the observed size differences could be variations in O-glycosylation. Prediction of theoretical O-glycosylation sites in the VNTR region revealed that CEL-WT has the highest number of potential sites (n=36), whereas DEL1, DEL4, DEL9 and DEL13 contain 13, 18, 23 and 26 sites, respectively (**Fig. 3**; Supporting Fig. 2). Thus, the proximal deletion variants DEL1 and DEL4 have fewer predicted O-glycosylation sites than the distal variants DEL9 and DEL13. This observation might at least to some degree explain the migration differences observed in Fig. 2.

**Figure 3.**
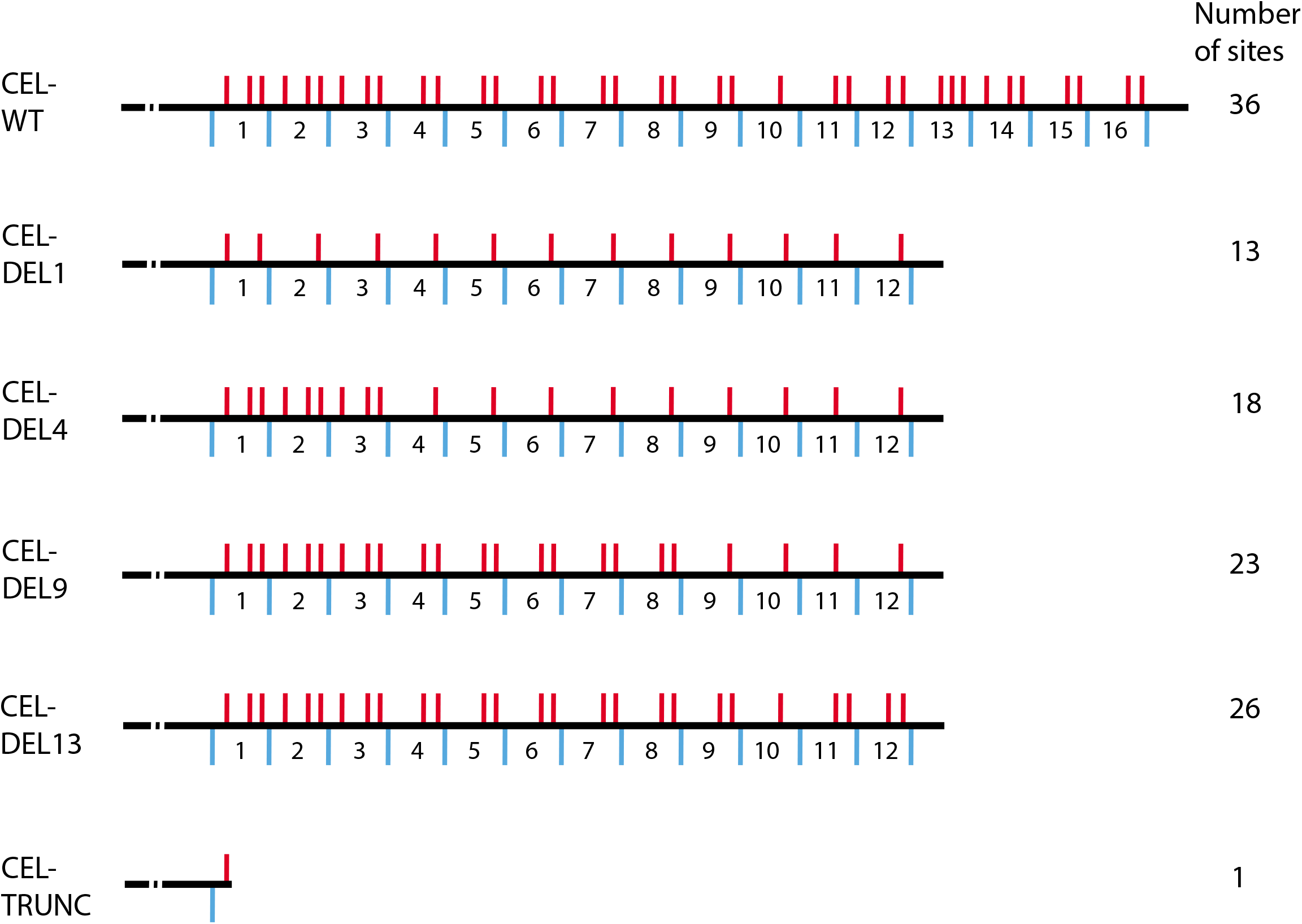
Potential O-glycosylation sites of the CEL VNTR region. For each CEL variant used in the study, theoretical O-glycosylation sites were predicted by the NetOGlyc3.1 server at http://www.cbs.dtu.dk/services/NetOGlyc/. These sites were all located in the VNTR region, and are indicated as red bars above the black line, with the total number of predicted sites listed on the right. VNTR segment numbers and borders (blue lines) of each variant are shown below the black line. For all deletion variants, the reading frame terminates within repeat 13.

To experimentally investigate the effect of O-glycosylation on the various DEL variants, an isogenic cell model (HEK293 cells) with and without knockout of the gene encoding the Cosmc chaperone was employed (25,26). As a result of the knockout, the enzymatic activity that provides O-glycan elongation beyond the initial GalNAc-α1 residue on O-linked glycoproteins (T-synthase) is absent in the so-called SimpleCells. We transfected SimpleCells and their non-altered HEK293 counterparts with the different CEL variants and analyzed the protein mass by western blotting (**Fig. 4**). We found that most variants showed a ~20 kDa reduction in band migration when expressed in SimpleCells, reflecting the presence of truncated O-glycans in the C-terminus of CEL. Furthermore, all CEL variants appeared to be less secreted into the medium of SimpleCells than in the medium of regular HEK293 cells. The only exception was TRUNC, which was secreted to a low but similar degree in both cell lines. Notably, DEL1 was the only variant whose migration corresponded to its predicted theoretical mass in the SimpleCells (~75 kDa). Moreover, while SimpleCells secreted CEL proteins with two different sizes into the medium, HEK293 cells secreted the variants as only one prominent band, representing the mature and heavily glycosylated form of the protein (compare the right panels of Fig. 4). However, proteins can undergo the initial O-glycosylation step in SimpleCells (i.e. attachment of GalNAcα1 onto Ser/Thr residues), which probably explains why size differences still are present after expressing the DEL variants in this cellular model. Taken together, our results indicate that full O-glycosylation is important for efficient secretion of CEL, and that the level of O-glycosylation varies considerably between the DEL variants.

**Figure 4.**
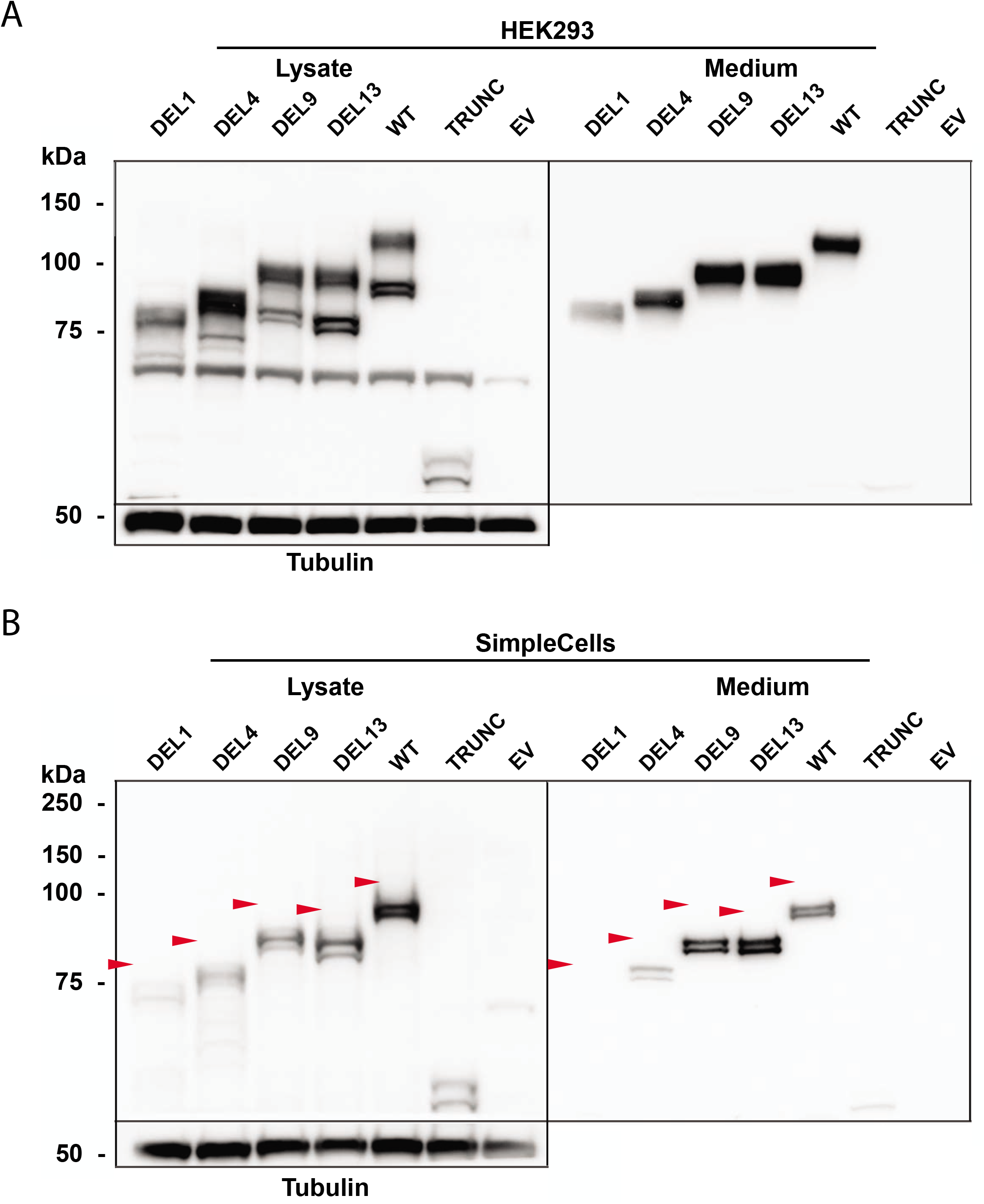
Expression of CEL variants in an O-glycosylation-deficient cellular model. HEK293 cells (*A*) and HEK293 cells with Cosmc knockout (SimpleCells) (*B*) were transiently transfected with plasmids encoding the CEL variants of Figure 1, and analyzed by western blotting. Cells transfected with empty vector (EV) were used as negative control. Tubulin expression was monitored as loading control. C-terminal O-glycosylation was assessed as shifts in protein migration in the SimpleCells when compared with expression in non-mutated HEK293 cells. The positions of the heaviest protein bands in each lane (except TRUNC), present in unmodified HEK293 cells but missing in the SimpleCells, are indicated in the lower image by red arrowheads. This illustrates the level of glycosylation in the CEL VNTR region. Secretion was evaluated based on loading equal amounts of protein from the conditioned media. One representative image of three independent experiments is shown.

### Expression of DEL1 and DEL4 in HEK293 cells increases ER stress

Expression of the DEL1 variant induces ER stress in both HEK293 cells and in the pancreatic acinar cell line AR42J (20). In the present study, we wanted to confirm this finding and to compare the ability of the different deletion variants to cause ER stress, both at the RNA and the protein level. In addition to analyzing an ER stress master regulator (the chaperone GRP78), all three branches of the unfolded protein response were investigated, namely the PERK, IRE1a and ATF6 pathways (27). HEK293 cells were transiently transfected with our set of CEL variants, followed by expression analysis of six marker genes (**Fig. 5**). Cells expressing DEL1 had significantly increased mRNA levels for *XBP1s* (*p* = 0.048), belonging to the IRE1a arm. For this variant, we also observed increased levels of borderline significance for *GRP78* and *ATF3* (*p* = 0.055 and *p* = 0.059, respectively). In addition, cells expressing the DEL4 variant showed higher levels of *GRP78* (*p* = 0.004), the ER folding chaperone *HYOU1* (*p* = 0.006) belonging to the ATF6 arm, as well as the pro-apoptotic factor *CHOP* (*p* = 0.047) of the PERK arm (Fig. 5). DEL9 and DEL13 did not result in significantly elevated levels of any of the tested marker genes.

**Figure 5.**
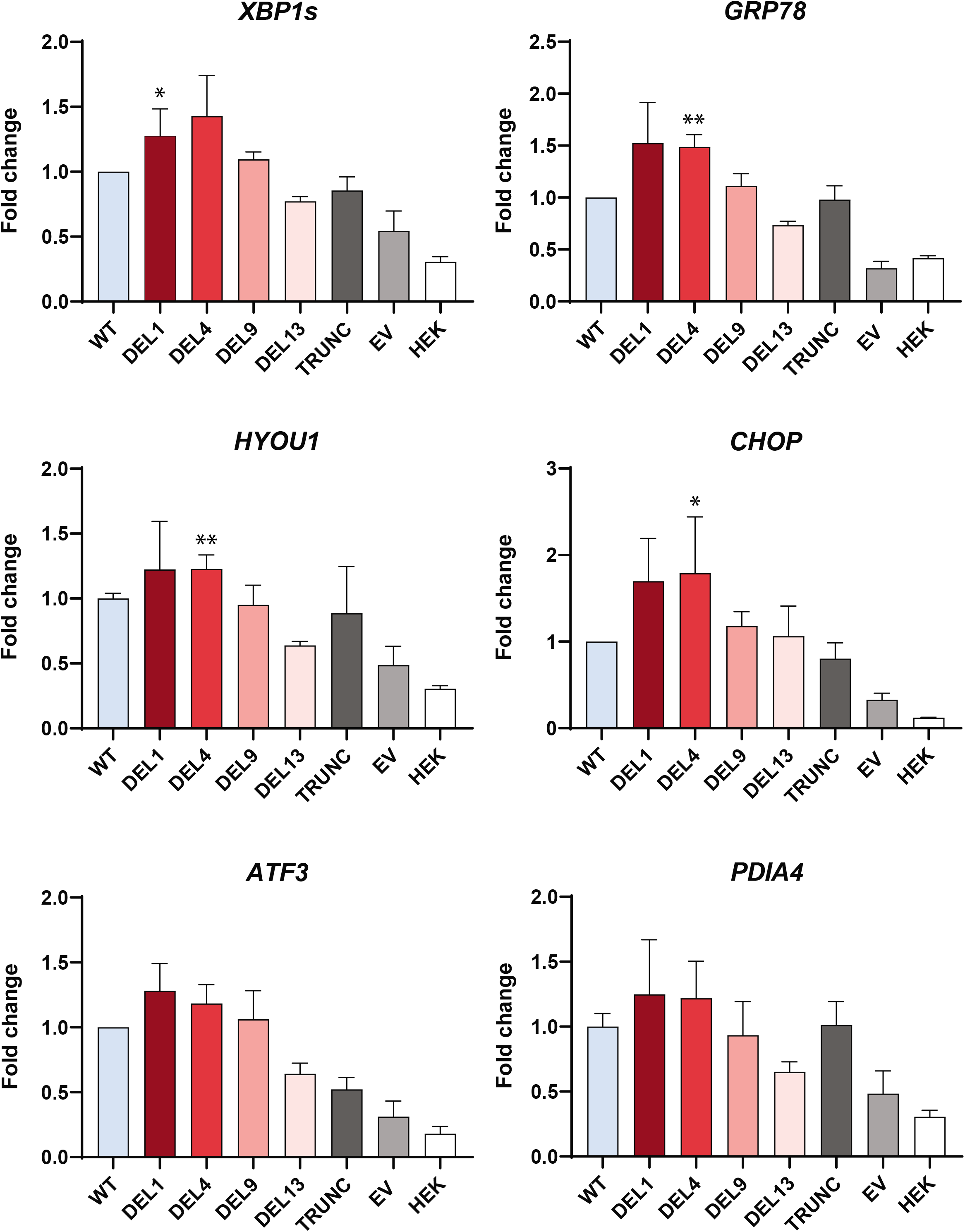
Effect of CEL variants on ER stress markers at the mRNA level. HEK293 cells were transiently transfected with plasmids encoding the CEL variants of Figure 1, and mRNA levels were analyzed by real-time quantitative PCR. Cells transfected with empty vector (EV) and untransfected cells (HEK) were used as negative controls. The mRNA expression levels were corrected to the geometric mean of three reference genes (*OAZ1, GAPDH, ACTB*) and then normalized to expression in CEL-WT transfected cells. Three independent experiments were performed. Error bars are standard deviation (SD). Statistical significance is indicated as follows: *, p < 0.05; **, p < 0.01.

At the protein level, we examined the effect of CEL variants on the ER chaperones GRP78 and calnexin, and the ER stress transducer PERK. Clear differences between the CEL variants were observed in the insoluble pellet fraction but not in the soluble lysate (**Fig. 6**; Supporting Fig. 3). Compared to WT, DEL1 significantly induced higher levels of GRP78 (*p* = 0.013), calnexin (*p* = 0.050) and PERK (*p* = 0.037) (Fig. 6). For DEL4 we observed a 210-fold increase in expression of the same ER stress markers although the differences did not quite reach statistical significance due to experimental variation (Fig. 6).

**Figure 6.**
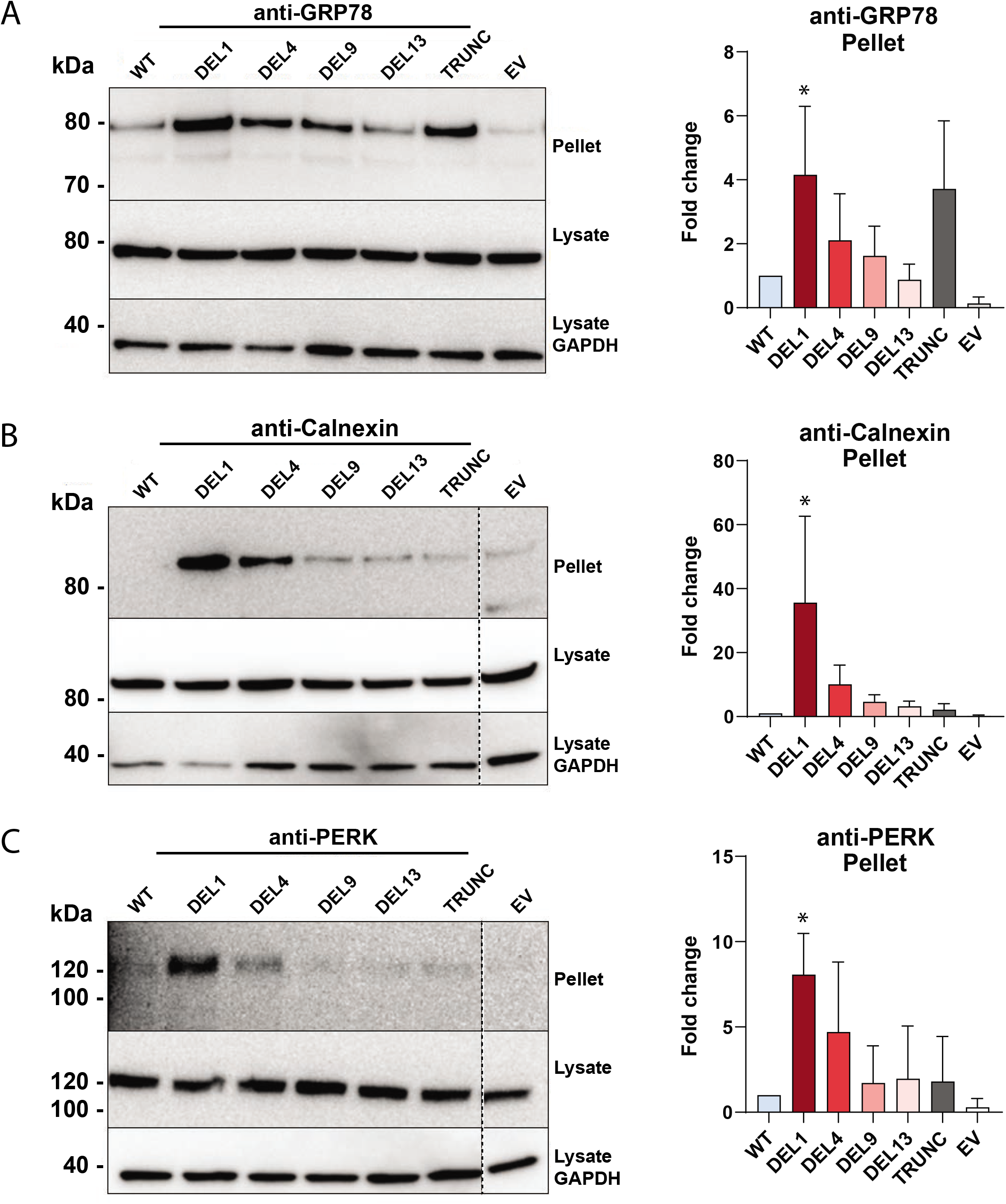
Effect of CEL variants on ER stress markers at the protein level. HEK293 cells were transiently transfected with plasmids encoding the CEL variants of Figure 1, and expression of ER stress markers was assessed by western blotting of the soluble (lysate) and insoluble (pellet) fraction. Cells transfected with empty vector (EV) were used as negative control. For each of the ER stress markers GRP78 (*A*), Calnexin (*B*), and PERK (*C*), one representative image of three independent experiments is presented. GAPDH expression was monitored for loading control. The right panels show quantification of band intensities for the insoluble pellet fraction after adjustment to the GAPDH levels and normalization to the expression in CEL-WT-transfected cells. Error bars are standard deviation (SD). Statistical significance is indicated as follows: *, p < 0.05. Stippled lines in *B* and *C* indicate unrelated lanes removed from the image.

When the data of Fig. 5 and Fig. 6 were considered together, the position of the deletion within the VNTR showed an impact on ER stress signaling: A stress response was induced by both DEL1 and DEL4, and this effect leveled off as the deletions were positioned more distally.

### Deletion variants display reduced lipolytic activity compared with normal CEL

To test whether the single-base deletions could impair the protein’s primary function, CEL enzymatic activity was measured. Conditioned medium from HEK293T cells expressing the CEL variants was collected, and the different variants were purified to near homogeneity as demonstrated by SDS-PAGE (**Fig. 7*A***) Lipolytic activity in the medium was then measured with trioctanoate as substrate (**Fig. 7*B***). To exclude that any changes in observed activity simply reflected altered levels of secretion, activities were first normalized against the molecular weight of the variant and then plotted after adjusting for the relative band intensities visualized in Fig. 7*A*. When compared with the activity of CEL-WT, the lowest activity was observed for the DEL1 protein (76% of WT activity, *p* < 0.001) (Fig. 7*B*). There was reduced activity also for DEL4 (84%, *p* = 0.002), DEL9 (90%, *p* = 0.008) and DEL13 (91%, *p* = 0.004). The TRUNC variant displayed the smallest reduction in enzyme activity (95%, *p* = 0.046), suggesting that a complete loss of the VNTR region does not affect lipolytic activity as much as an altered protein tail.

**Figure 7.**
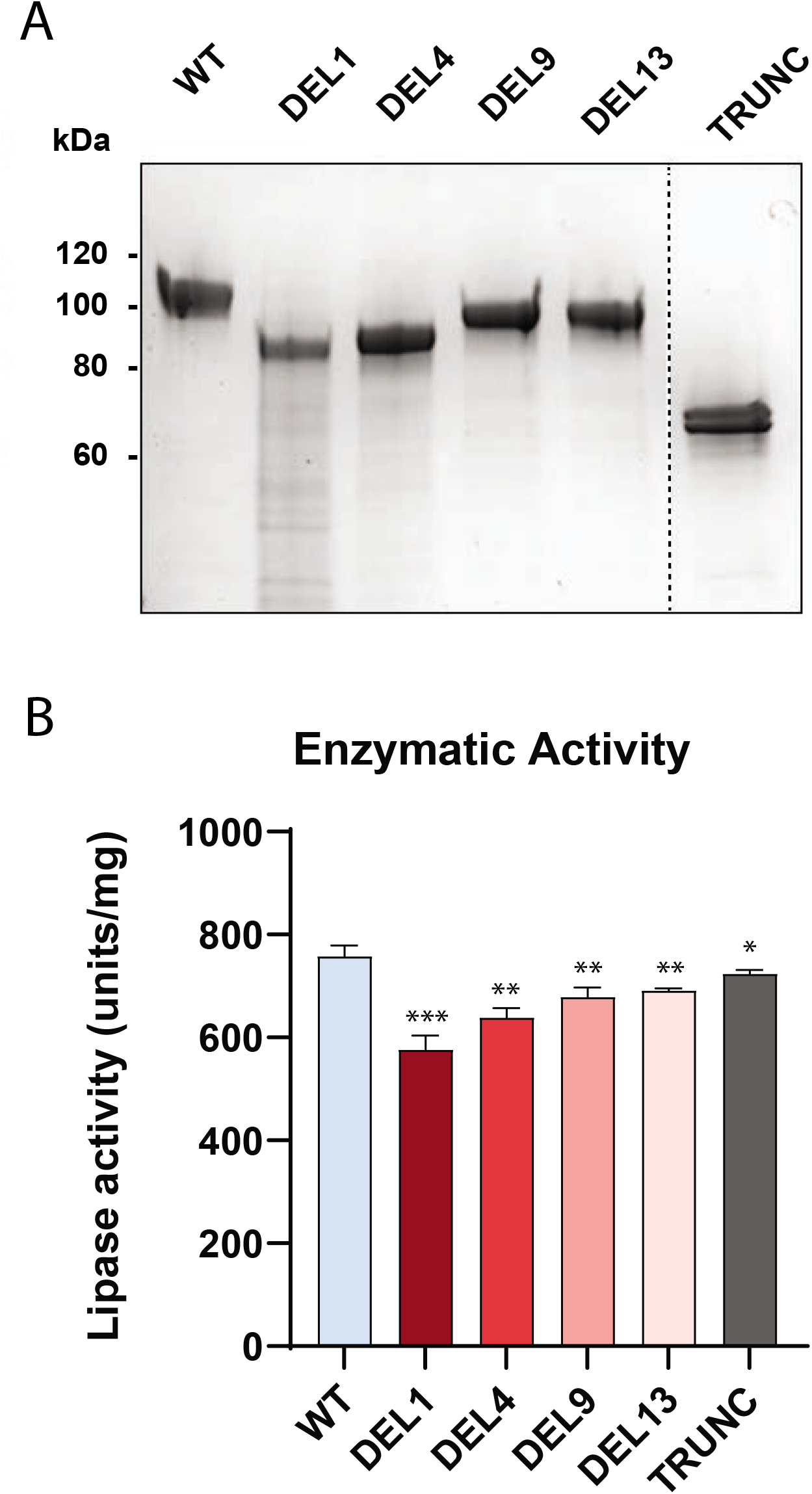
Enzymatic activity of the studied CEL variants. HEK293T cells were transiently transfected with plasmids encoding the CEL variants of Figure 1. *A*, SDS-PAGE of CEL variants purified from conditioned medium and stained with Gel-Code Blue. The same amount of protein was loaded in each lane. The stippled line indicates an unrelated lane removed from the image. One representative image of three independent experiments is shown. *B*, CEL lipolytic activity in the medium was measured with trioctanoate as substrate. The activity was adjusted for band density (from *A*) and molecular weight of each variant. Error bars are standard deviation (SD). Statistical significance is indicated as follows: *, p < 0.05; **, p < 0.01; ***, p < 0.001.

## DISCUSSION

The inherent variation and instability of VNTR sequences implies that these regions are excellent candidates for associations with human disease. We have previously investigated whether the number of VNTR segments in the *CEL* gene influence the risk for exocrine pancreatic disease; however, without finding any evidence that this is the case (9–11). In the current report, we have therefore focused on one-base pair deletions within the VNTR. These variants are extremely rare (5), and they are present in MODY8 families (16) or may occasionally be revealed when screening large cohorts of human DNA samples. As some cellular properties of the DEL1 variant are clearly altered from that of the normal CEL protein (16–20), we here investigated the impact of varying deletion positions within the VNTR.

Due to the frameshift effect, each of the four naturally occurring deletion variants analyzed in this study has a C-terminal amino acid sequence that differs substantially from that of normal CEL. Moreover, between the DEL proteins there is also considerable variation in the extension of normal versus aberrant sequence repeats (Fig. 1*B*). Discrepancies in cellular properties of the tested variants are therefore likely to reflect biochemical effects arising from their dissimilar amino acid compositions. Notably, the frameshifts alter the *pI* of the protein variants. The consequence is that the overall acidic charge changes to the basic side for DEL1, DEL4 and DEL9 (Fig. 1*B*). Another effect is the introduction of several cysteine residues: the aberrant tails of DEL1, DEL4 and DEL9 contain 12, 9 and 4 cysteines, respectively, whereas the tails of DEL13 and WT have none (Supporting Fig. 2). The positively charged VNTR region observed for DEL1 and DEL4, and to some degree for DEL9, could contribute to aggregation and pathogenicity through change of short- and long-range interactions between CEL protein monomers and between CEL and membrane surfaces (18). Important in this regard, is the observation of high-molecular weight bands only for DEL1 and DEL4 variants in the pellet fraction (Fig. 2*A*). The pronounced tendency to aggregate observed for DEL1 and DEL4 might also be attributed to formation of intra- and intermolecular disulfide bridges between opposing cysteine residues, as discussed in (20) and reported in other pathological proteins (28,29). DEL13 may be protected against aggregation because it keeps an overall acidic charge and has no cysteine residues in its VNTR region. Regarding the observed high-molecular weight forms of CEL, we speculate that they also could be protein complexes with binding partners such as the GRP94 chaperone (30) or ubiquitinated proteins targeted for proteasomal degradation.

The normal CEL VNTR region is heavily O-glycosylated, a post-translational modification considered important for proper secretion and stability of the protein (13,24). For several of the DEL variants, two or more bands were detected in the insoluble and soluble cell fractions (Fig. 2*A*), likely to represent fully glycosylated and partially glycosylated CEL, as previously observed (18). Although western blotting is not an accurate way of estimating the molecular mass of a protein, none of the tested variants migrated with a band size that corresponded to their unmodified amino acid composition. Notably, DEL1 and DEL4, both variants with dramatically fewer O-glycosylation sites, were found to be less well secreted (Fig. 2*A*).

The rationale behind expressing the DEL variants in HEK293 SimpleCells, a cellular model with deficient O-glycosylation, was to obtain variants with an even and truncated glycosylation profile. Ser and Thr residues in the VNTR region would then contain at most one GalNAc residue (with or without a terminal sialyl group). This might enable us to evaluate whether there was a substantial difference in the basic glycosylation pattern among the variants. When expressed in SimpleCells, the CEL proteins displayed a dramatic shift in band size. All variants migrated with a band size closer to that predicted from their amino acid composition, except for TRUNC, which has no VNTR sequence and only one theoretical site for O-glycosylation (Fig. 4*B*). Of note, TRUNC was the most abundant variant in the pellet fraction (Fig. 2). Also, when expressed in bacteria, truncated (and non-glycosylated) CEL was found as large aggregates recovered in inclusion bodies (31), further proving the importance of this post-translational modification on CEL secretion. The fact that the band sizes of the DEL variants still varied when expressed in SimpleCells despite very similar predicted molecular masses (Fig. 4) could indicate a large difference in glycan site occupancy. This might be due to the inability of the GalNAc-transferases (and sialidases) to reach or recognize certain areas in the aberrant tail, or a consequence of extensive intra- or intermolecular disulfide bonds.

The morphological changes of the pancreatic parenchyma observed in MODY8 are consistent with a phenotype of chronic pancreatitis (16,32). The classic model for understanding the pathogenesis of pancreatitis involves inappropriate intracellular activation of trypsinogen or failure of intracellular trypsin inactivation (33). However, it is becoming increasingly clear that there are also trypsin-independent disease mechanisms, caused by digestive-enzyme misfolding and ER stress (21,34–36). When investigating the effect of DEL variant expression on well-known ER stress markers at the RNA and protein level, there was a clear trend for DEL1 and DEL4 to induce ER stress markers, both when compared with WT and with the distal deletion variants DEL9 and DEL13 (Fig. 5). Increased signaling was detected in the three canonical branches of the unfolded protein response that are under the control of PERK, IRE1a and ATF6. This is consistent with ER stress caused by DEL1 and DEL4 protein aggregation, intra- and intermolecular disulfide bridge formation, reduced O-glycosylation and/or impaired transit through the secretory pathway. At the protein level, we found significant differences only in the insoluble pellet fraction (Fig. 6) and not in the soluble lysate (Supporting Fig. 3). For the folding chaperone GRP78, this could be explained by the protein binding tightly to misfolded CEL variants, thereby being pulled down in the pellet fraction as a protein complex.

All DEL variants displayed significantly decreased lipolytic activity compared with the normal CEL protein (Fig. 7). There was a gradual decrease in enzymatic function from DEL13 (91%) to DEL1 (76%), while TRUNC displayed an almost normal lipolytic activity (95%). The latter result is in line with a previous study showing that complete truncation of the VNTR region did not compromise the enzymatic ability of CEL (37,38). We find it unlikely that reduced lipolytic activities can explain pathogenicity of the DEL variants. Firstly, the observed activity differences do not appear large enough to alter physiological function of CEL, although the possibility remains that activity against other substrates than trioctanoate could be more severely affected. Secondly, a mouse model with a full-body knockout of CEL did not recapitulate any aspect of the MODY8 pancreatic phenotype (39). Thirdly, a recent study found that cytotoxic effects observed for different variants of the pathogenic CEL-HYB1 protein could not be attributed to differences in lipolytic activity (40,41). Finally, the presence of a normal gene in the heterozygous state is expected to compensate for any loss of activity from a rare *CEL* mutation. Taken together, all available data favor the hypothesis that pathogenic CEL variants initiate pancreatic disease through a dominant negative gain-of-function effect, rather than via a loss-of-function mechanism.

In summary, our study demonstrates that the position of single-base deletions in the *CEL* VNTR has great impact on protein cytotoxicity. When analyzed in transfected HEK293 cells, the pathogenic DEL1 and DEL4 variants found in MODY8 families had the strongest effect on protein secretion, protein aggregation and ER stress. On the other hand, DEL13 behaved more like normal CEL and must be considered a benign variant. We evaluate DEL9 to have intermediate effects but given that this variant so far has not been reported in any MODY8 family, we conclude that it is likely benign. The *CEL* VNTR is a region very prone to mutational events (5). Should single-base deletions in the second or third VNTR repeat (i.e. DEL2 and DEL3) be identified, we predict that they will be pathogenic. For any DEL5-DEL8 variant that might be observed, genetic segregation analysis and functional evaluation will be necessary to determine their pathogenic potential.

## EXPERIMENTAL PROCEDURES

### Plasmid constructions

Previously, we made the plasmid pcDNA3/CEL-WT 14R (with 14 VNTR segments) for expression of the normal CEL protein in mammalian cells (20). For this study, a *Psh*AI/*Xho*I cDNA fragment encoding normal CEL with 16 VNTR repeats was synthesized by Genscript and used to replace the corresponding segment in pcDNA3/CEL-WT 14R. All four *CEL* deletion constructs were then made through mutagenesis service provided by Genscript using the newly created pcDNA3/CEL-WT 16R as template. The CEL-TRUNC construct was generated by QuikChange II XL site-directed mutagenesis kit (Agilent Technologies) using pcDNA3/CEL-WT 14R as template (20).

For experiments comparing expression of epitope-tagged versus untagged CEL variants, the tagged plasmids have been described before (19). In short, the constructs were all based on a pcDNA3.1 vector containing a V5/His-tag (Invitrogen). Untagged constructs for comparison were prepared by introducing a stop codon into the pcDNA3.1/V5-His constructs using the same site-directed mutagenesis kit as above. The plasmid contains the *Xho*I restriction site directly before the epitope tag sequence, in which the stop codon was created.

### Antibodies and reagents

Two anti-CEL antibodies were used: a rabbit polyclonal antibody for immunoblotting (against the truncated CEL variant pV562Δ as described in (20)), and a mouse monoclonal antibody (As20.1) for immunocytochemistry (against the CEL globular domain), kindly provided by Prof. Olle Hernell (Department of Clinical Sciences, Umeå University, Umeå, Sweden). Additional primary antibodies were rabbit polyclonal anti-GRP78 antibody (Abcam, cat. 21685), rabbit monoclonal anti-calnexin (C5C9, cat. 2679) and mouse monoclonal anti-PERK (D11A8, cat. 5683) from Cell Signaling Technology, and mouse monoclonal anti-GAPDH (Santa Cruz Biotechnology, sc-47724). Secondary antibodies were HRP-goat anti-rabbit (Invitrogen, cat. 656120), HRP-donkey antimouse (Santa Cruz Biotechnology, sc-2318) and Alexa Fluor 488 anti-mouse (Invitrogen, cat. A11017).

RNeasy Mini Kit was from Qiagen. Polyethyleneimine (PEI) was from Polysciences. High Capacity cDNA Reverse Transcription Kit was from Applied Biosystems. Amersham Hybond P Western Blotting membranes (PVDF) was from GE Healthcare. TGX Precast protein gel (4–15%) and Blotting-Grade Blocker were from Bio-Rad. Poly-L-lysine, Fetal Bovine Serum (FBS), Phosphate Buffered Saline (PBS), Penicillin-Streptomycin (Pen-Strep) and cOmplete Mini Protease inhibitor cocktail tablets were from Sigma-Aldrich. ProLong Gold Antifade Solution containing DAPI nuclear stain, Antibiotic Antimycotic (AA) and Lipofectamine 2000 were from Invitrogen. RIPA Lysis Buffer (10X), Triton X-100 and Tween-20 were from Merck Millipore. NuPAGE Sample Reducing Agent (10X), NuPAGE LDS Sample Buffer (4X), NuPAGE Bis-Tris Protein gels, DMEM with high glucose (4500 mg/L), NuPAGE 10% Bis-Tris Protein gels, 1.0 mm, 10 well, NuPAGE 4-12% Bis-Tris Protein gels, 1.0 mm, 10 well, Pierce BCA Protein Assay, Pierce ECL Plus Western Blotting Substrate, Opti-MEM I Reduced Serum Medium and GelCode Blue stain reagent were obtained from Thermo Fisher Scientific.

### Cell culture of HEK293 and HEK293T cells

Human Embryonic Kidney 293 (HEK293) cells (Clontech Laboratories) were used for most experiments described in this study. For the enzymatic activity assay, HEK293T cells (ATCC) were used. HEK293 Cosmc-knock out cells (referred to as SimpleCells throughout the text) and their unaltered counterpart were obtained from the lab of Prof. Henrik Clausen (Copenhagen Center for Glycomics, Denmark). All cell lines were maintained in growth medium with high glucose (4500 mg/L) supplemented with 10% FBS and either 100 U/ml of AA or 10 mg/ml of Pen-Strep in humidified atmosphere with 5% CO_2_ at 37°C.

### Transient transfections

For transfection of HEK293 cells, Lipofectamine 2000 was used according to the manufacturer’s instructions. Cells were transfected with plasmids encoding CEL-WT, single-base deletion variants (DEL1, DEL4, DEL9, or DEL13) or CEL-TRUNC, either with or without the V5/His tag. Empty vector (EV; either pcDNA3 or pcDNA3.1-V5/His) was included as a negative control. Samples were harvested for analysis 48 h post-transfection for all experiments.

For the enzymatic activity assay, HEK293T cells were used. Cells were grown in 10-cm culture dishes in regular medium until reaching 70–90% confluence. Next, the cells were transfected with plasmids expressing different CEL variants using PEI. The PEI was dissolved in ddH_2_O at a concentration of 1 μg/μL, pH 7.4) and sterilized using a 0.22 μM filter. Prior to PEI transfection, cells were switched to 10 mL DMEM without FBS. For one dish, 15 μg plasmid DNA was mixed with 45 μL PEI solution in 500 μL Opti-MEM I Reduced Serum Medium. The mixture was added to the culture medium of HEK293T cells after 30 min incubation at room temperature. The cells were grown for 16-20 h before the medium was switched to 10 mL Opti-MEM I Reduced Serum Medium containing antibiotics. The cells were grown for another 24 h, the conditioned medium was harvested, and the cells were refreshed with another 10 mL of Opti-MEM I Reduced Serum Medium. The conditioned medium was again collected after 24 h incubation. The harvested media were pooled and filtered with 0.45 μm Steritop bottle top filters (Millipore) to remove debris prior to protein purification.

### Cell fractionation and collection of conditioned media

Conditioned medium was collected and analyzed as the medium fraction. The cells were lysed in ice-cold RIPA buffer supplemented with protease inhibitor. After 30 min incubation on ice, the lysate was centrifuged at 14,000 x g for 15 min at 4°C. The supernatant was isolated and analyzed as the soluble lysate fraction. The pellet was washed twice in PBS, boiled at 95°C in 2x LDS sample buffer and reducing agent and analyzed as the insoluble pellet fraction. All fractions were analyzed by western blotting.

### Western blotting

For detection of CEL protein, four μg of the soluble lysate fraction was employed. For the medium and insoluble pellet fractions, the volume loaded on the SDS gel was the same as for the corresponding lysate. All samples were added LDS sample buffer and reducing agent. Before loading, the medium and soluble lysate fractions were denatured on a heating block at 56°C for 15 min whereas the pellet fractions were denatured at 95°C for 15 min. The samples were loaded and separated by 10% SDS-PAGE at 90 V for 15 min, then at 180 V for 1 h. Proteins were transferred to a PVDF membrane by semi-wet blotting using XCell Blot Module chambers (Invitrogen) according to the manufacturer’s instructions. The proteins were detected using standard methods, with primary antibody incubation overnight at 4°C and secondary antibody incubation for 1 h at room temperature. Blots were developed using Pierce ECL Plus Western Blotting Substrate, and signals were analyzed using a Las 1000 Pro v2.6 software imager (Fujifilm). Band quantification was performed using the MultiGauge v3.0 software (Fujifilm). For ER stress markers, 20 μg (GRP78 and calnexin) or 40 μg (PERK) of the insoluble pellet fraction was loaded and separated by 4-12% SDS-PAGE. For ER stress analyses of the soluble lysate fractions, 10 μg protein was loaded on the gel. For analyses of CEL expression in transfected SimpleCells, 10 μg protein was loaded on the gel and separated by 10% SDS-PAGE.

### RNA extraction and real-time qPCR

Medium was removed and the cells were washed before total RNA was extracted using the RNeasy Mini Kit according to the manufacturer’s protocol. For cDNA synthesis, the High Capacity cDNA Reverse Transcription Kit was used with total RNA input of 20 ng for each cDNA synthesis reaction. The real-time PCR reaction was performed with a Rotor-Gene SyBR Green on a Rotor-Gene Q cycler (Qiagen). Gene expression was calculated as copies/μL using the standard curve approach. The standards were made with suitable primers in a conventional PCR reaction. *GAPDH*, *OAZ1* and *ACTB* were used as reference genes.

### Protein purification and enzymatic activity assay

CEL protein variants were purified by HiTrap Heparin HP Affinity Column (GE Healthcare Life Sciences) controlled by an ÄKTA pure protein purification system as described previously (20). Protein concentration was determined by measuring absorbance at 280 nm using the corresponding extinction coefficient for each CEL variant. Five μg of each purified protein was resolved by 4–15% TGX Precast protein gel and stained with GelCode Blue stain reagent to assess molecular mass, homogeneity and integrity. Functional characterization of recombinant CEL protein variants was carried out by measuring lipase activity against trioctanoate in the presence of 12 mM sodium cholate using a pH-stat (Radiometer Analytical) at pH 8.0. For each assay, 5 μg of purified CEL protein was used. The lipolytic activities were expressed in international lipase units/mg of enzyme (1 unit corresponds to 1 μmol of fatty acid released/min). The activities were adjusted according to densitometry and to the molecular weight of each variant.

### Immunocytochemistry and confocal microscopy

HEK293 cells were seeded onto coverslips (18 mm) coated with poly-L-lysine, transfected as described above and fixed in 3% paraformaldehyde for 30 min. Immunostaining was performed as previously described (17). Results were evaluated with an SP5 AOBS confocal microscope (Leica Microsystems) with 63x/1.4 NA NAHCX Plan-Apochromat oil immersion objective, ~1.2 airy unit pinhole aperture and appropriate filter combinations. Images were acquired with 405 diode and argon ion/argon krypton lasers (Leica) and processed using LAS AF lite (Leica), Photoshop CC and Adobe Illustrator CC (Adobe Systems).

### Statistical analysis

The two-tailed student’s t-test was employed for all statistical analyses. Statistical calculations were carried out using Microsoft Excel 2016. *P*-values < 0.05 were considered as statistically significant. Results are given as mean ± SD.

## ACKNOWLEDGEMENTS

The authors thank Olle Hernell, Umeå University, Sweden and Henrik Clausen, Copenhagen Center for Glycomics, University of Copenhagen, for kindly providing the As20.1 antibody and HEK293 SimpleCells, respectively.

## SUPPORTING INFORMATION

**Supporting Figure 1.**
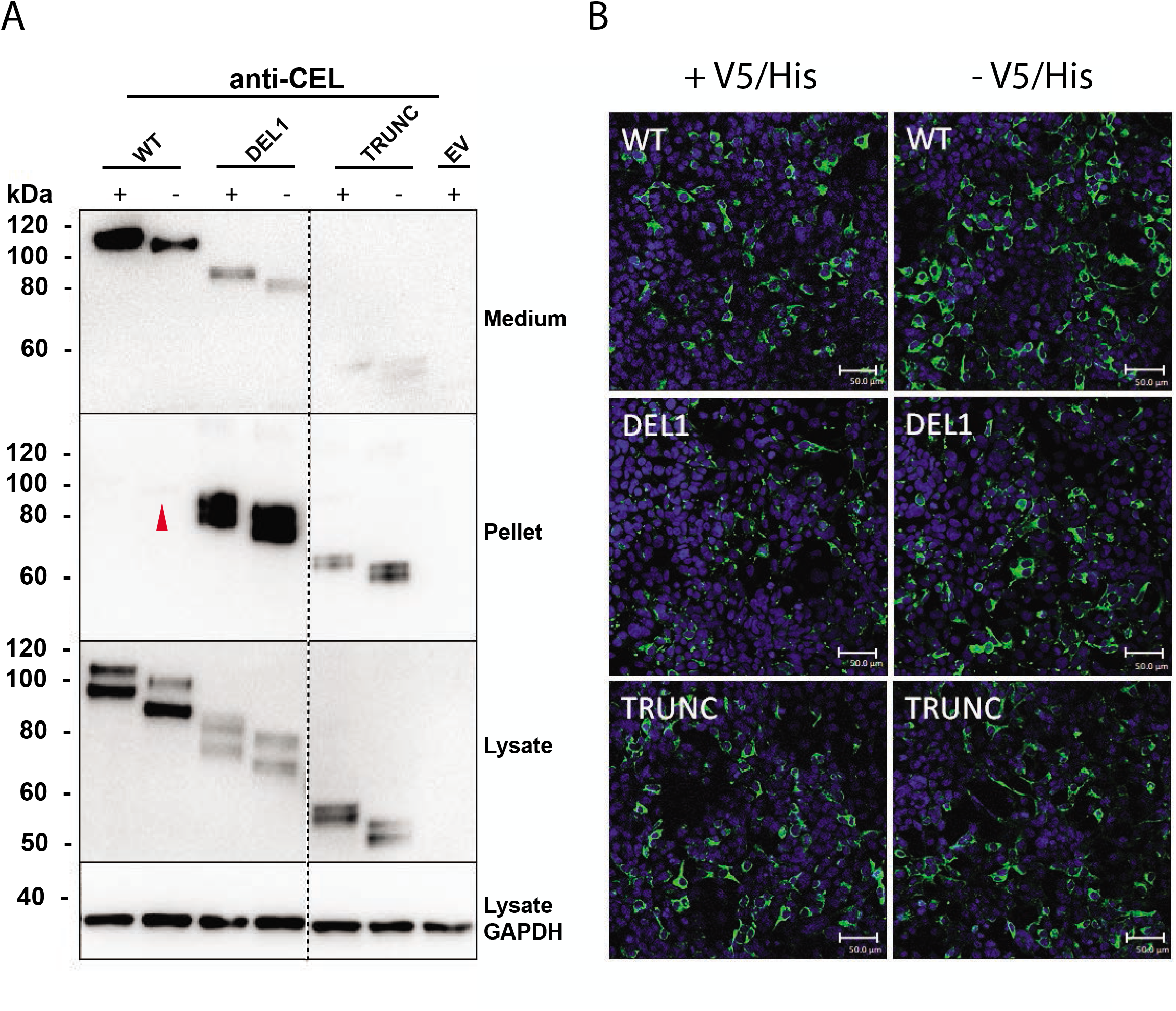
Secretion and intracellular distribution of tagged versus untagged CEL variants in HEK293 cells. HEK293 cells were transiently transfected with plasmids encoding CEL-WT, CEL-DEL1 or CEL-TRUNC with or without the V5/His tag (+/-). The expressed CEL proteins were examined by western blotting (*A*) and immunocytochemistry (*B*). *A*, secretion was evaluated as CEL levels detected in the conditioned medium. Intracellular distribution was assessed by analyzing the soluble (lysate) and insoluble (pellet) fractions of the cells. Cells transfected with empty vector (EV) were used as negative control, whereas GAPDH expression was monitored for loading control. The red arrow indicates a very weak band for untagged CEL-WT protein in the pellet fraction. One representative image of three independent experiment is shown. The stippled line indicates an unrelated lane removed from the images. *B*, transfected cells were subjected to immunostaining and confocal microscopy. The CEL protein variants are shown in green, while cell nuclei are shown in blue. Representative images of three independent experiments are presented. Scale bars are 50 μm.

**Supporting Figure 2.**
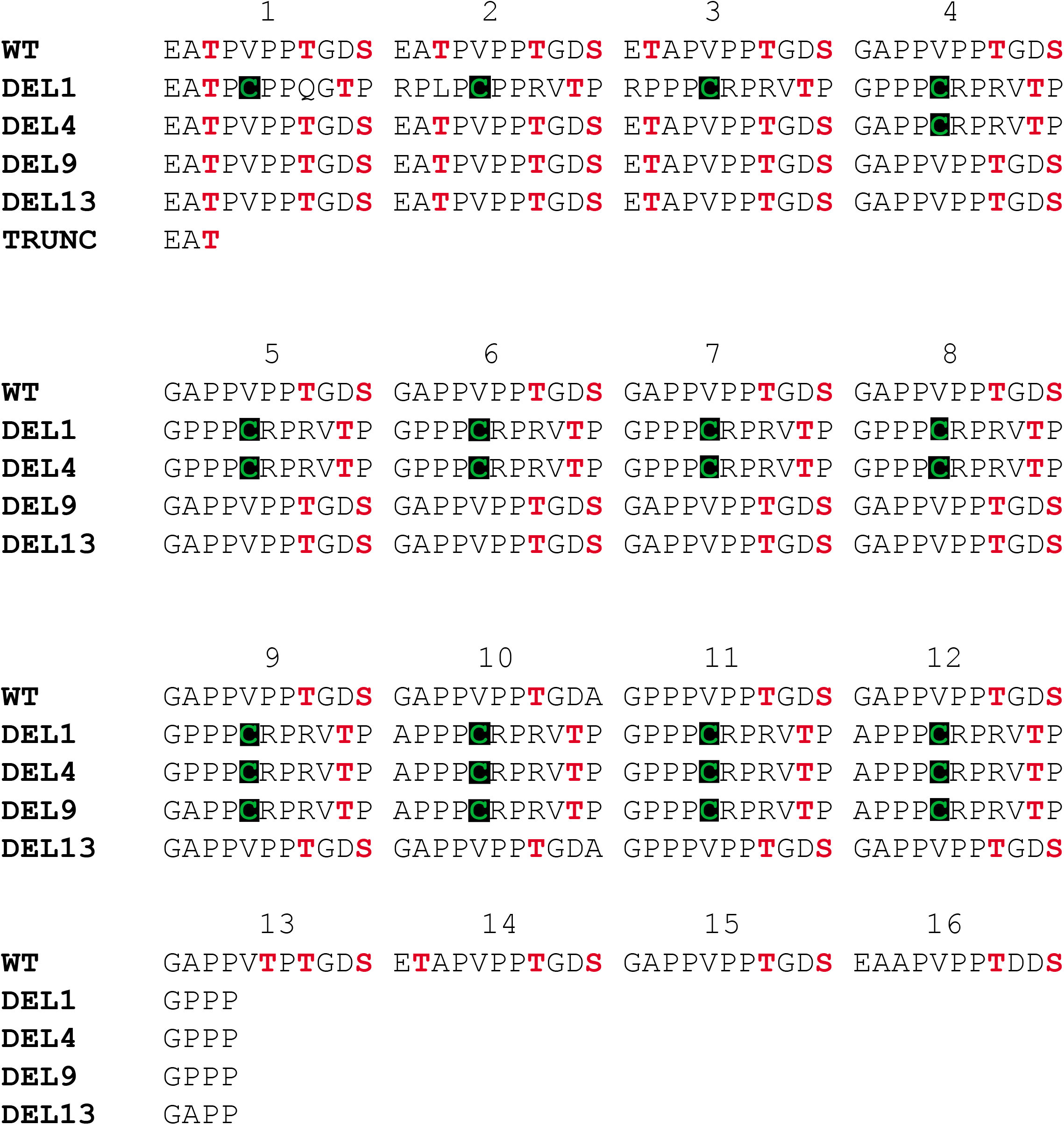
Amino acid sequences of the VNTR region of CEL-WT (16 repeats) and the investigated DEL variants. One-letter amino acid nomenclature is used, with predicted glycosylated residues (serine, S; threonine, T) highlighted in red. Cysteine residues (C), which may potentially form intra- and intermolecular disulfide bridges, are highlighted in green.

**Supporting Figure 3.**
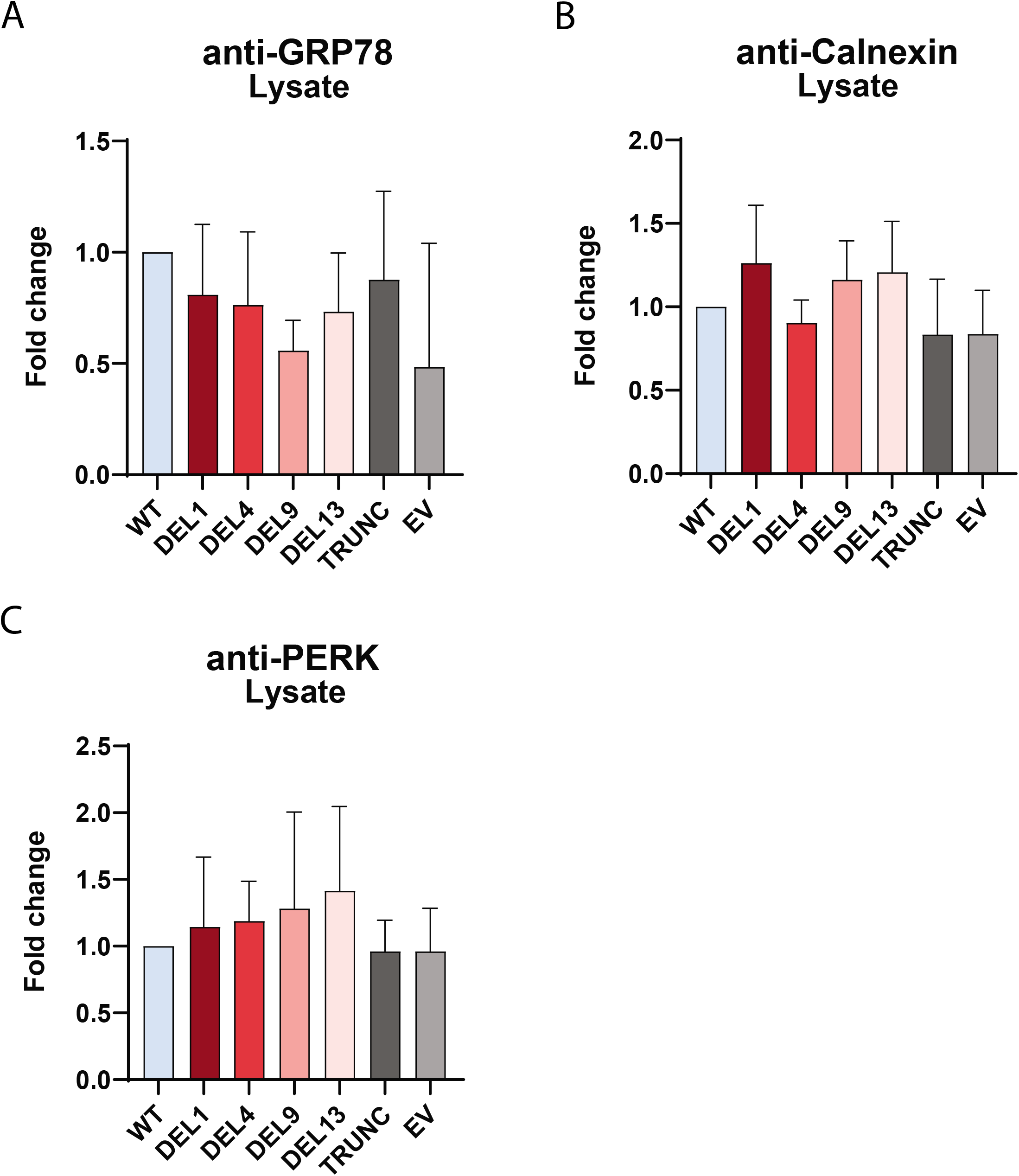
The effect of CEL variants on ER stress markers in the soluble lysate fraction of transfected HEK293 cells. This figure is based on western blots shown in Figure 6, and shows quantification of band intensities in the soluble lysate (three independent experiments) for each of the ER stress markers GRP78 (*A*), Calnexin (*B*), and PERK (*C*). Expression levels were first adjusted to GAPDH levels and then normalized to expression in CEL-WT-transfected cells. Error bars are standard deviation (SD).

